# Gal4 activation domain 9aaTAD could be inactivated by adjacent mini-inhibitory domain and reactivated by distal re-activation domain

**DOI:** 10.1101/110882

**Authors:** Martin Piskacek, Marek Havelka, Martina Rezacova, Andrea Knight

**Affiliations:** Laboratory of Cancer Biology and Genetic, Department of Pathological Physiology, Faculty of Medicine, Masaryk University Brno, Kamenice 5, 625 00 Brno, Czech Republic Laboratory; Gamma Delta T Cell Laboratory, Department of Pathological Physiology, Faculty of Medicine, Masaryk University Brno, Kamenice 5, 625 00 Brno, Czech Republic Laboratory

**Keywords:** Transcription, Gal4, 9aaTAD

## Abstract

The characterisation of the activation domains started three decades ago with Gcn4 and Gal4 activators. The amorphous character of the activation domains strongly hindered their definition. Moreover, during the attempts to localise the Gal4 activation domain, the artificial peptides, an unintended consequence of cloning, were responsible for artificial transcriptional activity of the several Gal4 constructs. These artefacts produced enormous experimental bias and misconception. The presence of inhibitory domains in some Gal4 constructs made the misperception even worse. Previously, we reported that the nine amino acid transactivation domain, 9aaTAD, is the exclusive activation domain in the Gal4 protein. The activation domain 9aaTAD could be identified in Gal4 paralogs Oaf1, Pip2, Pdr1, Pdr3 and other activators p53, E2A and MLL. Surprisingly, the activation domain 9aaTAD was reported as misconception for Gal4 activator. Here we demonstrated that small region of 10 amino acids adjacent to the Gal4 activation domain 9aaTAD is an inhibitory domain, which the authors included in their constructs. Moreover, we identified Gal4 region, which was able to the reverse the inhibitory effect. The 9aaTAD re-activation domain was localized to the 13 amino acid long region. In this report we clarified the numerous confusions and rebutted supposed 9aaTAD misconception.

**Summary:** The activation domain 9aaTAD has decisive function in Gal4 activation. Gal4 activation domain 9aaTAD could be inhibited by adjacent region of 10 amino acids. The inhibited Gal4 activation domain 9aaTAD could be reactivated by 13 amino acid long Gal4 region. The activation domains 9aaTAD could be identified by our 9aaTAD prediction algorithm, especially in the Gal4 family.

## Introduction

The Nine amino acid Transactivation Domain, 9aaTAD, is universally recognized by the transcriptional machinery in eukaryotes (1–3). Currently, the 9aaTAD family comprises of over 40 members including Gal4, Oaf1, Pip2, Pdr1, Pdr3, Leu3, Tea1, Pho4, Gln3, Gcn4, Msn2, Msn4, Rtg3, E2A, MLL, p53-TADI, p53-TADII, FOXO3, NF-kB, NFAT, CEBPA/E, ESX, ELF3, ETV1, KLF2/4, EBNA2, VP16, HSF1, HSF2, HsfA, Gli3, Sox18, PIF, Dreb2a, MTF1, OREB1, WRKY45, NS1, MKL1, VP16, EBNA2, KBP220, ECapLL, P201, AH, and B42 transcription factors. We and others have shown that the activation domains 9aaTAD have competence to activate transcription as small peptides (1–16). We have established previously the 9aaTAD prediction service online (www.piskacek.org). The curated 9aaTAD activation domains had been annotated on protein database UniProt (www.uniprot.org/9aaTAD).

The Gal4 transcription factor was model eukaryotic activator used in the early 90s to study gene activation. In the Mark Ptashne lab, great efforts had been made to determine Gal4 activation domains. Multiple activation domains were reported in series of studies. Nevertheless, we had found that all previously reported activation domains in the Gal4 have none considerable transactivation potential and are not involved in activation of transcription. We demonstrated that artificial peptides included in the original Gal4 constructs (17) and fusion regions (18) were responsible for the artificial transcriptional activities in numerous reported Gal4 constructs (19). All these artefacts produced enormous bias and misconception.

From the Gal4 ortholog study on the transcription factor Oaf1, we have reported new concept for activation domain. The activation domain 9aaTAD, we had identified in Oaf1, enabled us to identified further activation domain in the Gal4 family. We recognized the activation domain 9aaTAD in the Gal4 activator (857-871 aa) and experimentally proved it's ability to activate transcription (2). The Gal4 activation domain 9aaTAD location did not correspond to any of the previously defined activation domains AD-I, AD-II or AD-III from Ptashne lab, but rather correlate with the Gal80 binding region and with the region deleted in Gal4 loss-of-function mutant (1-852 aa)(2).

Strikingly, strong arguments for the 9aaTAD misconception were reported from Hahn lab, Warfield et al., 2014, PNAS, edited by M.Ptashne (20). According to the authors, the Gal4 9aaTAD domain is none functional activation domain and the 9aaTAD algorithm did not predict activation function. The 9aaTAD misconception was further worsen and disseminated by review Erkina and Erkine et al., 2016, Epigenetics & Chromatin (21).

In this study, we revised the 9aaTAD misconception for the Gal4 activator (20, 21). We identified a 9aaTAD adjacent inhibitory domain, which strongly reduced the 9aaTAD function. Because our experimental data well correlated with results from Hahn lab, we could declare experimental conformity. The authors included in their 9aaTAD constructs the adjacent Gal4 region, which cause the inhibition of the 9aaTAD activation domain. For unknown reason, the authors did not included in their experiments a construct with the original length of the activation domain 9aaTAD, reported by us previously. Such a set of constructs would immediately indicate to the authors a presence of the inhibitory domain in their longer 9aaTAD construct. Strikingly and very concerning, the positive control used by authors (20), the construct with extraordinary strong and well defined Gal4 activation domain from the Ptashne lab (18), was inactive in their system (Fig. 1).

**Fig 1.**
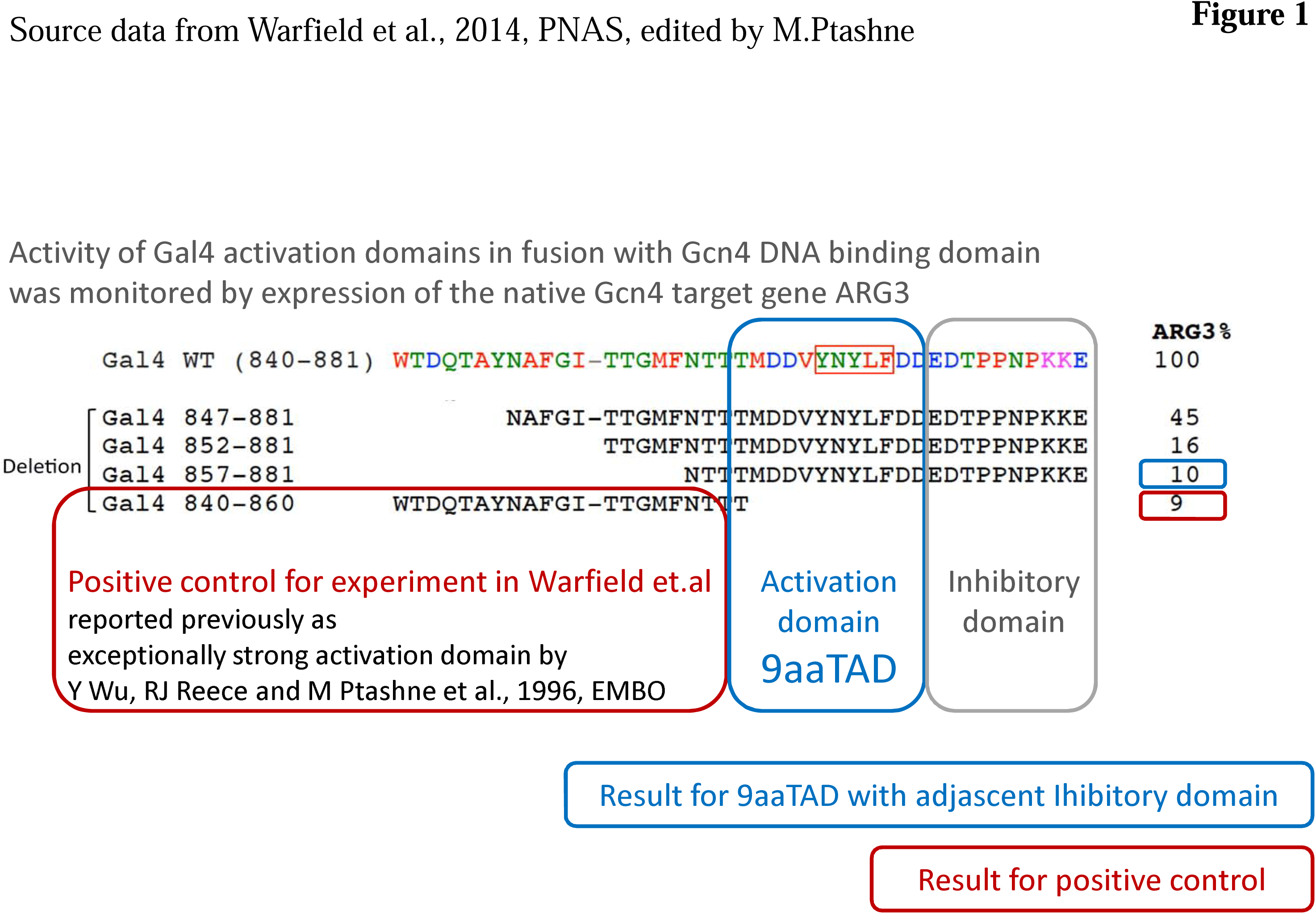
Head to head assayed activity of the Gal4 activation domains. The Gal4 - Gcn4 DNA binding hybrid were assayed for activation of transcription (Source data from Warfield et al., 2014, PNAS, edited by M.Ptashne). The red labelled extraordinary strong activation domain well defined by Ptashne lab (Gal4 840-860 aa), which served in their experiment as positive control and ii) the blue labelled activation domain 9aaTAD, defined by our lab (Gal4 857-870 aa) with grey labelled adjacent amino acids identified as inhibitory domain (871-881). The author conclude from this result that unlike red labelled activation domain defined by Ptashne lab (Gal4 840-860 aa), the blue labelled activation domain 9aaTAD reported by us previously is not activation domain.

## Materials and Methods

### Constructs

The construct pBTM116-HA was generated by an insertion of the HA cassette into the *Eco*RI site of the vector pBTM116 (HA cassette nucleotide sequence: TGG CTG - GAATTA - GCC ACC ATG GCT TAC CCA TAC GAT GTT CCA GAT TAC GCT GTC GAG ATA - GAATTC, which render in amino acids sequence: W L - E L - A T M A Y P Y D V P D Y A V E I - E F). The constructs G1-G45 and H1-H45 were generated by PCR and subcloned into pBTM116 *Eco*RI and *Bam*HI sites. All construct have a spacer of three amino acids inserted into the *Eco*RI site; peptide -NNN- (NNN cassette: GAATTC - AATAATAAT, which render in peptide: EF - NNN). All constructs were sequenced by Eurofins Genomics. Further detailed information about constructs, primer sequences are available on the request.

### Assessment of enzyme activities

The β-galactosidase activity was determined in the yeast strain L40 (4, 22). The strain L40 has integrated the *lacZ* reporter driven by the *lexA* operator. In all hybrid assays, we used 2μ vector pBTM116 for generation of the LexA hybrids. The yeast strain L40, the Saccharomyces cerevisiae Genotype*:MATa ade2 his3 leu2 trp1 LYS::lexA-HIS3 URA3::lexA-LacZ*, is deposited at ATCC (#MYA-3332). For β-galactosidase assays, overnight cultures propagated in YPD medium (1% yeast extract, 2% bactopeptone, 2% glucose) were diluted to an A_600_ of 0.3 and further cultivated for two hours and collected by centrifugation. The crude extracts were prepared by vortexing with glass beads for 3 minutes. The assay was done with 10 ul crude extract in 1ml of 100 mM phosphate buffer pH7 with 10 mM KCl, 1 mM MgSO4 and 0.2% 2-Mercaptoethanol; reaction was started by 200 ul 0,4% ONPG and stopped by 500 ul 1 M CaCO3. The average value of the β-galactosidase activities from three independent experiments is presented as a percentage of the reference with the standard deviation (means and plusmn; SD; n = 3). We standardized all results to previously reported Gal4 construct HaY including merely the activation domain 9aaTAD with the activity set to 100% (3).

### Western Blot Analysis

The crude cell extracts were prepared in a buffer containing 200 mM Tris-HCl, pH 8.0, 1 mM EDTA, 10% glycerol (v/v), separated by SDS-PAGE, and blotted to nitrocellulose. The immuno-detection of proteins was carried out using mouse anti-HA antibody (#26183, ThermoFisher Sci) or mouse anti-LexA (#306-719, EMD Millipore Corp). The secondary antibodies used were anti-mouse IgG antibodies conjugated with horserAD-Ish peroxidase (#A9044, Sigma Aldrich). The proteins were visualized using Pierce ECL (#32106, ThermoFisher Sci) according to the manufacturer’s instructions.

## Results

### Inhibition of the Gal4 9aaTAD by adjacent region

In the study Warfield et al., 2014, PNAS, edited by Mark Ptashne (20), the ability of two Gal4 activation domains were assayed and evaluated head-to-head: i) the extraordinary strong activation domain defined by Ptashne lab (*Gal4 840-860* aa)(18), which served in their experiment as positive control and ii) the new-coming activation domain 9aaTAD, defined by our lab (Gal4 857-870 aa) (1, 2). Their data showed that both constructs are almost inactive (Figure 1).

From our recent study (19), we know well why their positive control, the activation domain defined by Ptashne lab, could not work in their experiments as a strong activator - reported errors in Ptashne constructs (3, 19).

What we did not know, why their longer 9aaTAD construct with adjacent Gal4 region was inactive. Initially, we inspected the Gal4 constructs reported previously and identified a construct reported from Ptashne lab (17), which included merely the activation domain 9aaTAD with the same adjacent Gal4 region (*construct ID5-3S, Gal4 DNA binding domain 1-74 aa fused with C-terminal Gal4 region 851-881 aa; 0,05% activity of the construct pMA210 with full length sequence of the Gal4 protein,* (17))(Fig. 2). Also this construct was inactive, what indicate real experimental observation reported in Warfield et al. (20). Therefore, we supposed that the function of the 9aaTAD activation domain was inhibited by the Gal4 adjacent region in both constructs with the same inhibitory domain. The different hybrid systems used by Ptashne or Hahn lab could be therefore excluded for artificial inhibitory effects and rather natural Gal4 phenome could be expected (*Gal4 DNA binding domain and Gcn4 DNA binding domain hybrids*).

**Fig. 2.**
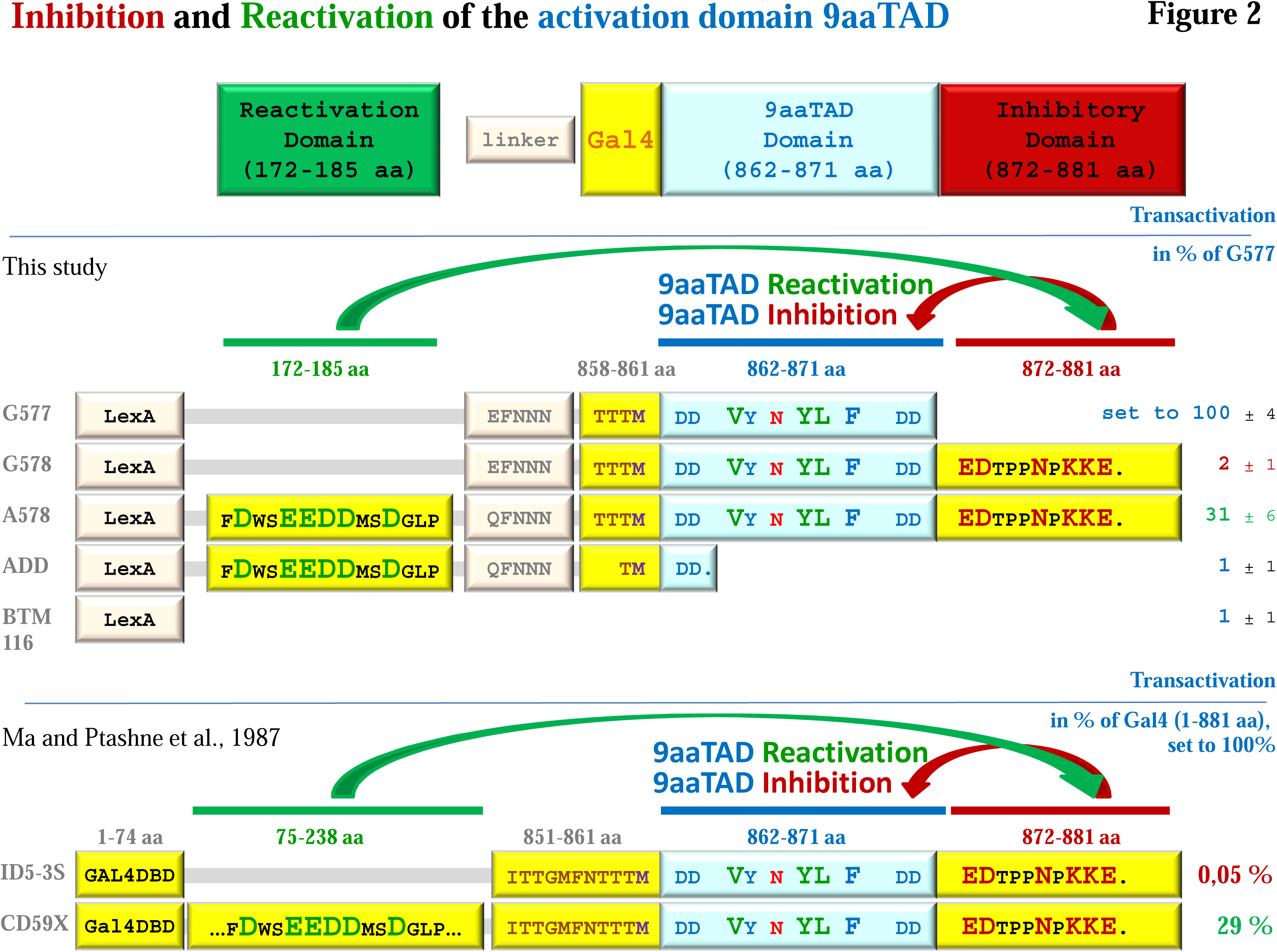
Inhibition and Re-activation of the activation domain 9aaTAD in the Gal4. To distinguish activity of the activation domain 9aaTAD with and without adjacent and distal regions, a set of the Gal4 constructs were generated and tested. The Gal4 - LexA hybrid constructs (BTM116 backbone) were assayed in L40 strain for activation of transcription. The average value of the β-galactosidase activities from three independent experiments is presented as a percentage of the reference with standard deviation (means and plusmn; SD; n = 3). We standardized all results to Gal4 construct G577 including merely the activation domain 9aaTAD with the activity set to 100%. The LexA is *E.coli* DNA binding domain generally used for the generation of hybrid constructs for transactivation assay. The regions of Gal4 protein in the constructs are noted and graphically presented. Single dot means end of protein sequence. We showed graphically a similar result reported previously by others, which is in well agreement with our results and significantly force our findings (Ma and Ptashne et al., 1987).

Noteworthy, the presence of inhibitory domains in Gal4 constructs are anymore surprising observations, because multiple inhibitory domains were already reported for the Gal4 protein (23) and could be expected in the inactive Gal4 constructs.

To test expected inhibition of the 9aaTAD activation domain by the adjacent region, we assayed the 9aaTAD activation domain in LexA hybrid assay with and without the adjacent region (*LexA DNA binding domain hybrids with the GAL4 activation domain 9aaTAD*). We have generated LexA construct G577 with the 9aaTAD activation domain (*Gal4 857-871 aa, see protein expression in (19)) and LexA construct G578 with adjacent Gal4 region according to the authors (Gal4 857-881 aa, see protein expression in* (19)). The activation functions of the LexA-Gal4 hybrids were monitored in L40 strain (*2 μ vector pBTM116 was used for all constructs; strain L40 with integrated lacZ reporter driven by lexA operator was used to assay the activation of transcription*). The construct G578 with the activation domain 9aaTAD and adjacent Gal4 region was completely inactive, but the construct G577 (*including merely the activation domain 9aaTAD*) was strong activator, which activity exceed the activity of the Gal4 terminal constructs (19). The sequence difference of analogous constructs determined presence of inhibitory domain in the Gal4 adjacent region (Fig. 2). Thus, our and authors results are in a good agreement (*experimental conformity*), just the presence of obligatory controls (*positive control, which is not inactive in conducted experiment and which could activate transcription*) and the conclusions made (*logical conclusions*) largely differ.

### Re-activation of the inhibited activation domain 9aaTAD

The 9aaTAD re-activation could be observed in the Gal4 construct CD59X, which is 500 times more active as the shorter construct ID5-3S (*CD59X: Gal4 DNA binding domain 1-74 aa, with adjacent Gal4 region 75-238 aa and C-terminal Gal4 region 851-881 aa; 29% activity of the construct pMA210 with full length sequence of the Gal4 protein; ID5-3S: Gal4 DNA binding domain 1-74 aa fused with C-terminal Gal4 region 851-881 aa; 0,05% activity of the construct pMA210 with full length sequence of the Gal4 protein,* (17)).

From this could be concluded that the N-terminal region was responsible for re-activation, what we have further followed. This observation also reinforced presence of inhibitory domain in the original construct use by Warfield et al., 2014. Furthermore, the full length sequence of the Gal4 protein, which includes both activation domain 9aaTAD and adjacent inhibitory region is constitutively active (in the L40 strain, where inhibitory protein Gal80 is deleted) and strongly indicated the activation domain 9aaTAD re-activation. Therefore, there must be a region in the Gal4 protein, which interfere with the inhibitory domain and provided release of the activation domain 9aaTAD.

Because of the strong charging of both 9aaTAD activation domain and the C-terminal inhibitory domain, we have first tested N-terminal acidic region of the Gal4 protein for the 9aaTAD re-activation function. We have generated construct A578 by insertion of a cassette coding for FDWSEEDDMSDGLP peptide (*Gal4 region 172-185 aa*) in to the G578 construct, where the 9aaTAD domain function was inhibited by adjacent region. The construct A578 showed about 30% activity of the construct G577 including the activation domain 9aaTAD without adjacent region (*positive control*). The negative control (*construct ADD*) generated by insertion of the cassette coding for FDWSEEDDMSDGLP peptide in to construct missing the 9aaTAD domain, did not activate transcription at all (Fig. 2).

With further investigations, we could identified another Gal4 region with ability to interfere with the 9aaTAD adjacent inhibitory domain. The 9aaTAD re-activation could be observed in the construct ID-3, which is 300 times more active as the shorter construct ID5-3S *(ID5-3: Gal4 DNA binding domain 1-74 aa fused with C-terminal Gal4 region 768-881 aa; 18% activity of the construct pMA210 with full length sequence of the Gal4 protein; ID5-3S: Gal4 DNA binding domain 1-74 aa fused with C-terminal Gal4 region 851-881 aa; 0,05% activity of the construct pMA210 with full length sequence of the Gal4 protein,* (17)).

From above we have localized 9aaTAD re-activation regions I-long (*Gal4 region 148-236 aa, 500 times reactivation in the construct CD59X,* (17)), I-short (*Gal4 region 172-185 aa, 30 times reactivation in the construct A578, this study*). II-long (*Gal4 region 768-857 aa, 300 times reactivation in the constructs ID5-3,* (17)) and II-short (*Gal4 region 840-857 aa, 10 times reactivation in the construct Gal4 (840-881 aa),* (20), Fig. 1).

The re-activation region II-short fully overlap with sequence of the reported construct Gal4 (*840-881 aa*) from Hahn lab (*Warfield et al., 2014*) (20), which is 10 times more active as their shorter Gal4 construct (*857-881 aa*). This explained full activity of all Gal4 C-terminal constructs (*857-881 aa*) .n any hybrid systems (LexA DBD used by us, Gal4 DBD used by Ptashne or Gcn4 DBD used by Hahn).

### Gal4 replicas are activation domains 9aaTAD

Previous study reported G80BP-A and G80BP-B peptides, which were selected from peptide-library by interaction with inhibitory protein Gal80 (24). Surprisingly, these Gal4 replicas were functional transactivation domains, which directly bind to mediator MED15 as original Gal4 TAD. The authors recognized well the structural and functional relationship between G80BP-A and G80BP-B peptides and the native domains. Nevertheless, the G80BP-A and G80BP-B peptides did not share significant similarity with the Gal4 activation domain (25).

We expect that both G80BP-A and G80BP-B peptides are solely Gal4 9aaTAD replicas selected in original screen by binding cavity of the inhibitory protein Gal80 (Fig. 3). The fact, that there are any significant sequence similarity G80BP-A and G80BP-B peptides with Gal4 reflect promiscuous pattern of the 9aaTAD family. In both G80BP-A and G80BP-B peptides, we have found putative activation domains 9aaTAD by our online 9aaTAD prediction. We generated LexA DNA binding domain constructs with the predicted nine amino acid long domains, which we considered as 9aaTAD replicas.

**Fig. 3.**
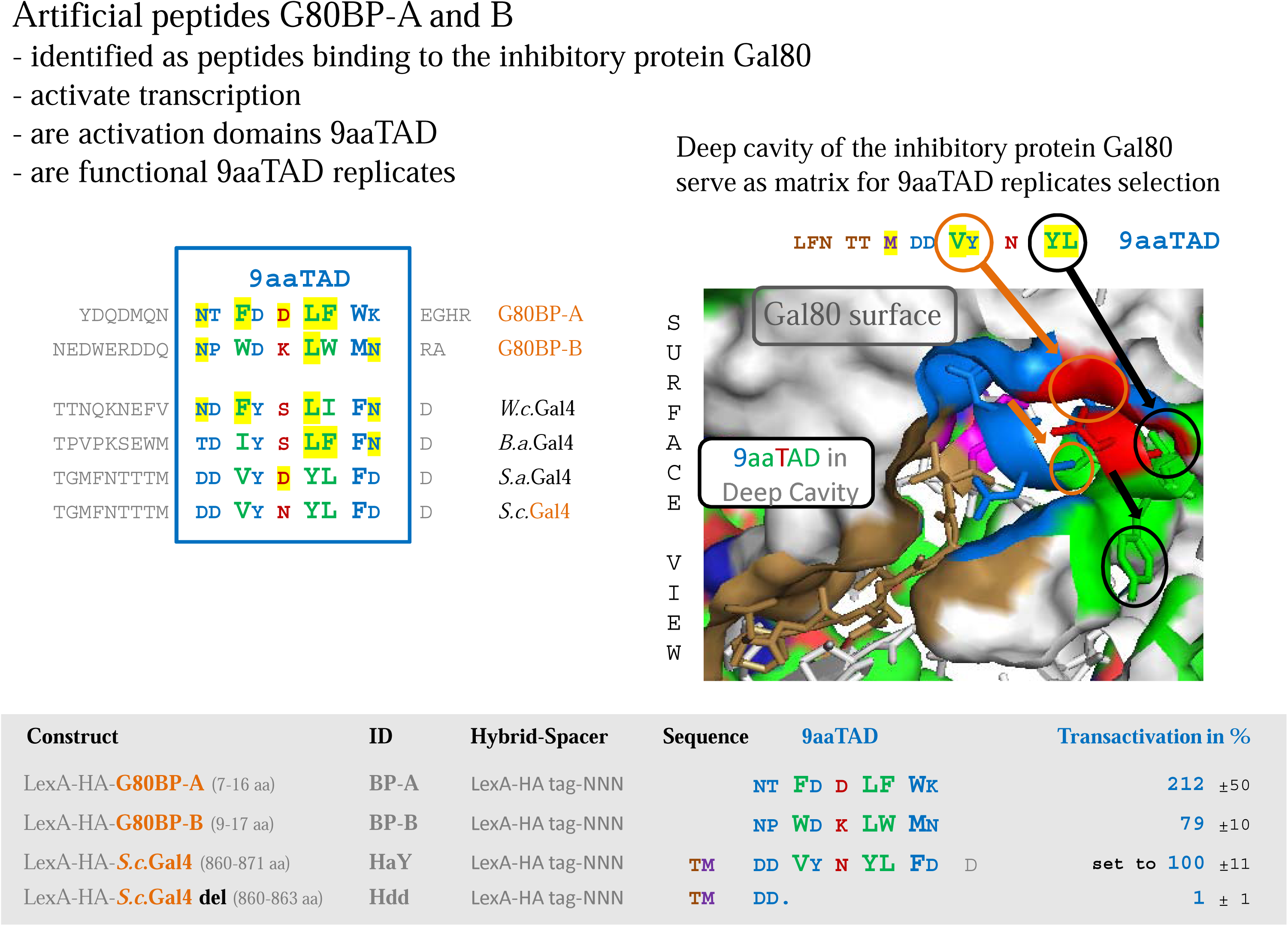
Gal4 replicas are activation domains 9aaTAD. The LexA - Gal4 hybrid constructs (BTM116 backbone) were assayed in L40 strain for activation of transcription. To test predicted activation domains 9aaTAD in the G80BP-A and G80BP-B peptides for activation potential, LexA - Gal4 hybrids constructs with predicted sequences were generated. The average value of the β-galactosidase activities from three independent experiments is presented as a percentage of the reference with standard deviation (means and plusmn; SD; n = 3). We standardized all results to Gal4 construct HaY including merely the activation domain 9aaTAD with the activity set to 100%. The LexA is *E.coli* DNA binding domain generally used for the generation of hybrid constructs for transactivation assay. The binding cavity of the inhibitory protein Gal80 specifically bind major determinant of the activation domain 9aaTAD, what explained selection of 9aaTAD replicas in reported screen, the G80BP-A and G80BP-B peptides. The Gal80 protein surface binding cavity and Gal4 peptide with activation domain 9aaTAD is shown.

The resulted activities of the predicted activation domains 9aaTAD were comparable with the Gal4 activation domain 9aaTAD (Fig. 3). Collectively, both Gal4 replicas are activation domains 9aaTAD, which are transcriptionally active. The G80BP-A and G80BP-B Gal4 replicas were assigned to the 9aaTAD family.

### 9aaTAD domain prediction and conservation in the Gal4 family

Recently reported, the 9aaTAD prediction algorithm is highly redundant nine-residue AD signatures and does not predict AD function in Gal4 transcription factor (20, 21). Although, any prediction has limited accuracy and use, the 9aaTAD prediction made by our algorithm for the Gal4 activator well correlated with our experimental results. The online 9aaTAD prediction for the *S.c.* Gal4 protein revealed two hits, where one is in the DNA binding domain (false hit) and the second in the C-terminal region, which is well know activation domain (Fig. 4). The 9aaTAD prediction is useful for the Gal4 orthologs. The Gal4 orthologs were identified by ExPASy SIB BLAST. We found that the Gal4 activation domain 9aaTAD is conserved in the family (Fig. 5). For the most distal Gal4 ortholog identified by us, so far, yeast *Wickerhamomyces ciferrii, W.c.* Gal4 (K0KTZ9_WICCF), we revealed a single prediction hit from the online 9aaTAD prediction. Noteworthy, the *S.c.*Gal4 and *W.c.*Gal4 have no homology in the C-terminal region (Fig. 5). Moreover, the prediction algorithm was reported in 2007 and the genome result for *Wickerhamomyces ciferrii* genome was released in 2012, what demonstrated none related design of prediction algorithm and target sequences (26). We generated LexA DNA binding domain construct HaW with the predicted *W.c.* Gal4 activation domain 9aaTAD (*766-778 aa*). The activation functions of the LexA-Gal4 hybrids were monitored in L40 strain (*2 μ vector pBTM116 was used for all constructs; strain L40 with integrated lacZ reporter driven by lexA operator was used to assay the activation of transcription*). The *W.c.*Gal4 9aaTAD strongly activated transcription and confirmed thus our correct 9aaTAD prediction (Fig. 6).

**Fig 4.**
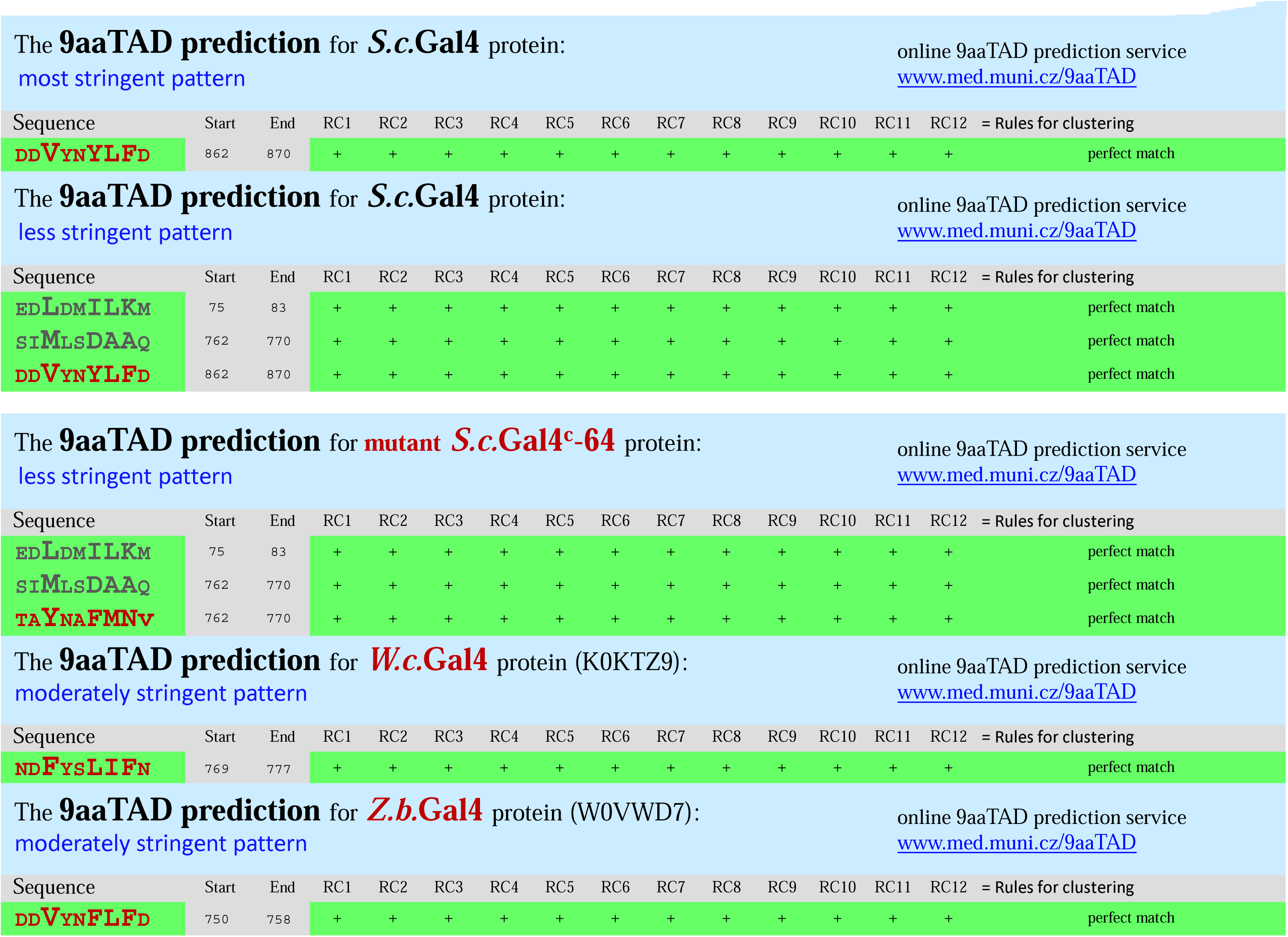
Prediction of the activation domains in Gal4 proteins by 9aaTAD algorithm. The 9aaTAD prediction was done online (http://www.piskcek.org). The false hits are in grey (according experimental data for Gal4, Piskacek et al. 2017) and patterns used are noted (less and moderate stringent).

**Fig 5.**
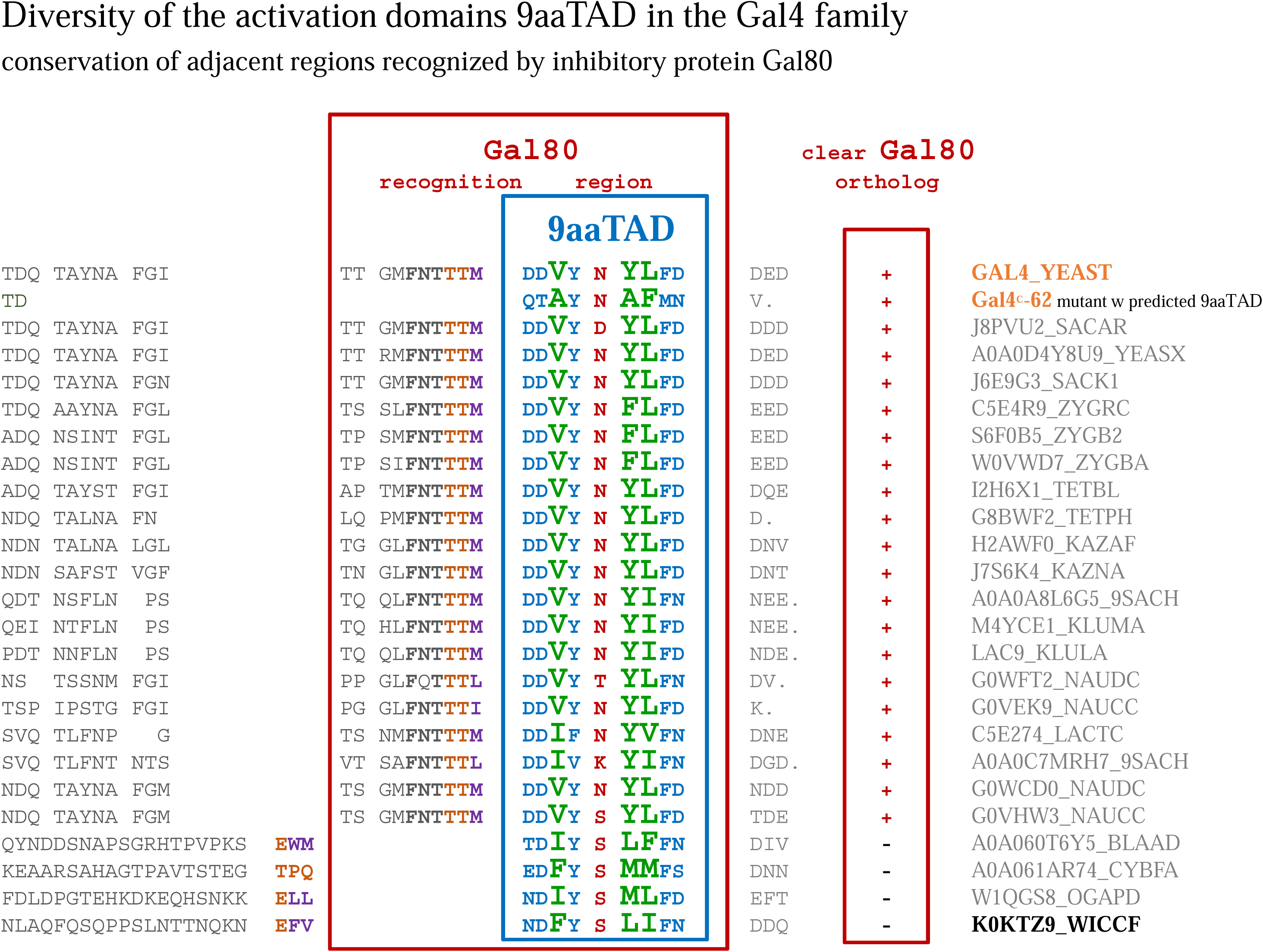
Alignment of the predicted Gal4 activation domains. The Gal4 orthologs alignment of the activation domains 9aaTAD and adjacent regions recognized by inhibitory protein Gal80. The presences of the clear Gal80 homologs, which correspond with conservation of the 9aaTAD adjacent regions, are marked. The sequence of transcriptional constitutively active Gal4^c^-62 mutant (independent on Gal80), which included predicted activation domain, is shown.

**Fig 6.**
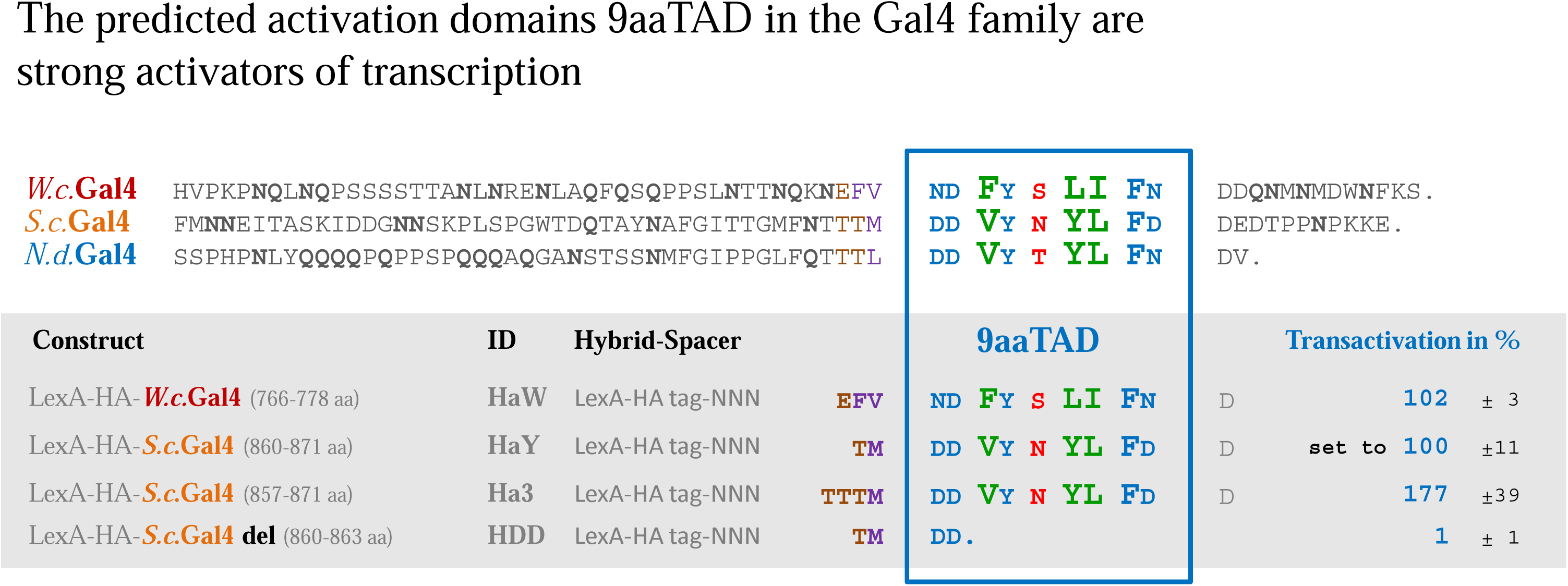
Predicted 9aaTAD in the Gal4 ortholog was tested for activation of transcription. To test predicted activation domain 9aaTAD for activation potential, LexA - Gal4 hybrids construct with predicted sequences were generated and tested. The LexA - Gal4 hybrid constructs (BTM116 backbone) were assayed in L40 strain for activation of transcription. The average value of the β-galactosidase activities from three independent experiments is presented as a percentage of the reference with standard deviation (means and plusmn; SD; n = 3). We standardized all results to Gal4 construct HaY including merely the activation domain 9aaTAD with the activity set to 100%. The LexA is *E.coli* DNA binding domain generally used for the generation of hybrid constructs for transactivation assay. The C-terminal sequence of selected Gal4 ortholog is shown to oversee homology in activation domain region, Gal4 from *Saccharomyces cerevisiae* (this study), *Wickerhamomyces ciferrii* (the most distal clear ortholog we could identified), *Naumovozyma dairenensis* (the example with short C-terminus, high identity for activation domain 9aaTAD and region recognized by Gal80 with *S.c.* Gal4, but low similarity in the rest of C-terminal region).

## Discussion

This report is in part a revision of the supposed 9aaTAD misconception (20, 21) and reported inhibitory and reactivation mechanism of activation domain 9aaTAD in Gal4 activator. Technical errors, which generated the Gal4 dogmas and controversy, are clarified here.

The Gal4 Dogma #1 stood up for activation domains AD-I and AD-II and their acidic character (Ma and Ptashne et al. 1987, Cell) (17). We had revised competence of the activation domains AD-I and AD-II to activate transcription. We found that both activation domains have none considerable transactivation potential and are not involved in activation of transcription (19). The artificial peptides included in the original Gal4 constructs, an unintended consequence of cloning, were responsible for the artificial transcriptional activity.

The Gal4 Dogma #2 adhered on the strong activation domain AD-III (Gal4 840-857 aa, construct pRJR200)(3). Already the position of activation domain AD-III generated serious concern. Firstly, this activation domain was located outside of both previously reported activation domains AD-I and AD-II (18). Furthermore, unlike the activation domain 9aaTAD, the AD-III position was outside of the region recognized by inhibitory protein Gal80 (27). We had revised reported experimental data and found that also this activation domain AD-III has none considerable transactivation potential and could not be involved in activation of transcription (19). We had found that the part of the Gal4 DNA binding domain (*region 92-100 aa*) fused to part of the activation domain AD-III (*region 840-843 aa*) in the original construct from Ptashne lab generated strong artificial activation domain, another unintended consequence of cloning [19].

The recent Gal4 Dogma #3 came from Hahn lab. The authors suppose to found evidence for the 9aaTAD misconception, Warfield et al., 2014, PNAS, edited by Mark Ptashne (20) ("Highly redundant nine-residue AD signatures", "has little activity, demonstrating that the proposed nine-residue motif does not predict AD function"). The 9aaTAD misconception was further worsen and disseminated by review Erkina and Erkine et al., 2016, Epigenetics & Chromatin (21).

The authors included adjacent Gal4 region in their 9aaTAD constructs, which cause the inhibition of the 9aaTAD activation domain. This another unintended consequence of cloning in the Gal4 history added "odd bit" to the activation domain 9aaTAD, which have strong impact on the activation function. Nevertheless, their 9aaTAD construct, including the adjacent inhibitory region, still activated transcription more than their positive construct with extraordinary strong and well defined activation domain from the Ptashne lab.

In agreement with authors, we also found that the 9aaTAD constructs with the adjacent region was inactivated. We identified inhibitory domain in the Gal4 C-terminal region, what explained our, their and others observations for various 9aaTAD constructs with and without adjacent region (17, 20). The inhibition of the activation domain 9aaTAD had produced enormous bias and misconception in recent (20, 21) but also in the very earlier Gal4 studies (17). Collectively, we could explained rationally all inconsistency in reported experiments by Warfield et al., 2014 (20) and rebut supposed 9aaTAD misconception.

The activation domain 9aaTAD is well conserved in the Gal4 family. The 9aaTAD algorithm recognized activation domains in *S.c.* Gal4 and other Gal4 orthologs and paralogs (2, 4). The 9aaTAD prediction is useful for identification of some but unfortunately not for all activation domains. This study revised the reported 9aaTAD dogmas, rebutted 9aaTAD misconceptions, identified experimental errors, clarified confusions and justified the function of the activation domains 9aaTAD in the Gal4 proteins.

## Competing interests

The author declares no competing interests.

## Funding

This work was supported by the Ministry of Health of the Czech Republic 15-32935A.

## References

1. Piskacek M, Vasku A, Hajek R, Knight A (2015) Shared structural features of the 9aaTAD family in complex with CBP. Mol Biosyst 11 (3): 844–851.

2. Piskacek S, et al. (2007) Nine-amino-acid transactivation domain: establishment and prediction utilities. Genomics 89(6): 756–768.

3. Piskacek M, Havelka M, Rezacova M, Knight A (2016) The 9aaTAD Transactivation Domains: From Gal4 to p53. PLoS ONE 11(9):e0162842.

4. Baumgartner U, Hamilton B, Piskacek M, Ruis H, Rottensteiner H (1999) Functional analysis of the Zn(2)Cys(6) transcription factors Oaf1p and Pip2p. Different roles in fatty acid induction of beta-oxidation in Saccharomyces cerevisiae. J Biol Chem 274(32): 22208–22216.

5. Sandholzer J, Hoeth M, Piskacek M, Mayer H, de Martin R (2007) A novel 9-amino-acid transactivation domain in the C-terminal part of Sox18. Biochem Biophys Res Commun 360(2): 370–374.

6. Lindert U, Cramer M, Meuli M, Georgiev O, Schaffner W (2009) Metal-responsive transcription factor 1 (MTF-1) activity is regulated by a nonconventional nuclear localization signal and a metal-responsive transactivation domain. Mol Cell Biol 29(23): 6283–6293.

7. Piskacek M (2009) 9aaTAD Prediction result (2006). Nature Precedings. doi:10.1038/npre.2009.3984.1.

8. Piskacek M (2009) Common Transactivation Motif 9aaTAD recruits multiple general co-activators TAF9, MED15, CBP and p300. Nature Precedings. doi:10.1038/npre.2009.3488.2.

9. Piskacek M (2009) 9aaTADs mimic DNA to interact with a pseudo-DNA Binding Domain KIX of Med15 (Molecular Chameleons). Nature Precedings. doi:10.1038/npre.2009.3939.1.

10. Hong JY, et al. (2011) Phosphorylation-mediated regulation of a rice ABA responsive element binding factor. Phytochemistry 72(1): 27–36.

11. Shekhawat UKS, Ganapathi TR, Srinivas L (2011) Cloning and characterization of a novel stress-responsive WRKY transcription factor gene (MusaWRKY71) from Musa spp. cv. Karibale Monthan (ABB group) using transformed banana cells. Mol Biol Rep 38(6): 4023–4035.

12. Lou S, et al. (2012) Human parvovirus B19 DNA replication induces a DNA damage response that is dispensable for cell cycle arrest at phase G2/M. J Virol 86(19): 10748–10758.

13. Matsushita A, et al. (2012) The nuclear ubiquitin proteasome degradation affects WRKY45 function in the rice defense program. Plant J. doi:10.1111/tpj. 12035.

14. Aguilar X, et al. (2014) Interaction studies of the human and Arabidopsis thaliana Med25-ACID proteins with the herpes simplex virus VP16- and plant-specific Dreb2a transcription factors. PLoS ONE 9(5):e98575.

15. Scharenberg MA, et al. (2014) TGF-β-induced differentiation into myofibroblasts involves specific regulation of two MKL1 isoforms. J Cell Sci 127(Pt 5):1079–1091.

16. Qiu Y, et al. (2015) HEMERA Couples the Proteolysis and Transcriptional Activity of PHYTOCHROME INTERACTING FACTORs in Arabidopsis Photomorphogenesis. Plant Cell. doi:10.1105/tpc.114.136093.

17. Ma J, Ptashne M (1987) Deletion analysis of GAL4 defines two transcriptional activating segments. Cell 48(5): 847–853.

18. Wu Y, Reece RJ, Ptashne M (1996) Quantitation of putative activator-target affinities predicts transcriptional activating potentials. EMBO J 15(15): 3951–3963.

19. Piskacek M, Havelka M, Rezacova M, Knight A (2017) The 9aaTAD Is Exclusive Activation Domain in Gal4. PLoS ONE 12(1):e0169261.

20. Warfield L, Tuttle LM, Pacheco D, Klevit RE, Hahn S (2014) A sequence-specific transcription activator motif and powerful synthetic variants that bind Mediator using a fuzzy protein interface. Proc Natl Acad Sci USA 111(34):E3506–3513.

21. Erkina TY, Erkine AM (2016) Nucleosome distortion as a possible mechanism of transcription activation domain function. Epigenetics Chromatin 9:40.

22. Miller JH (1972) Experiments in molecular genetics (Cold Spring Harbor Laboratory).

23. Stone G, Sadowski I (1993) GAL4 is regulated by a glucose-responsive functional domain. EMBO J 12(4): 1375–1385.

24. Han Y, Kodadek T (2000) Peptides selected to bind the Gal80 repressor are potent transcriptional activation domains in yeast. J Biol Chem 275(20): 14979–14984.

25. Højfeldt JW, Van Dyke AR, Mapp AK (2011) Transforming ligands into transcriptional regulators: building blocks for bifunctional molecules. Chem Soc Rev 40(8): 4286–4294.

26. Schneider J, et al. (2012) Draft genome sequence of Wickerhamomyces ciferrii NRRL Y-1031 F-60-10. Eukaryotic Cell 11(12): 1582–1583.

27. Kumar PR, Yu Y, Sternglanz R, Johnston SA, Joshua-Tor L (2008) NADP regulates the yeast GAL induction system. Science 319(5866): 1090–1092.

